# Do forest over- and understory respond to the same environmental variables when viewed at the taxonomic and trait level?

**DOI:** 10.1101/2021.09.06.459058

**Authors:** Kenny Helsen, Yeng-Chen Shen, Tsung-Yi Lin, Chien-Fan Chen, Chu-Mei Huang, Ching-Feng Li, David Zelený

## Abstract

While the relative importance of climate filtering is known to be higher for woody species assemblages than herbaceous assemblage, it remains largely unexplored whether this pattern is also reflected between the woody overstory and herbaceous understory of forests. While climatic variation will be more buffered by the tree layer, the understory might also respond more to small-scale soil variation, next to experiencing additional environmental filtering due to the overstory’s effects on light and litter quality. For (sub)tropical forests, the understory often contains a high proportion of fern and lycophyte species, for which environmental filtering is even less well understood. We explored the proportional importance of climate proxies and soil variation on the species, functional trait and (functional) diversity patterns of both the forest overstory and fern and lycophyte understory along an elevational gradient from 850 to 2100 m a.s.l. in northern Taiwan. We selected nine functional traits expected to respond to soil nutrient or climatic stress for this study and furthermore verified whether they were positively related across vegetation layers, as expected when driven by similar environmental drivers. We found that climate was a proportionally more important predictor than soil for the species composition of both vegetation layers and trait composition of the understory. The stronger than expected proportional effect of climate for the understory was likely due to fern and lycophytes’ higher vulnerability to drought, while the high importance of soil for the overstory seemed driven by deciduous species. The environmental drivers affected different response traits in both vegetation layers, however, which together with additional overstory effects on understory traits, resulted in a strong disconnection of community-level trait values across layers. Interestingly, species and functional diversity patterns could be almost exclusively explained by climate effects for both vegetational layers, with the exception of understory species richness. This study illustrates that environmental filtering can differentially affect species, trait and diversity patterns and can be highly divergent for forest overstory and understory vegetation, and should consequently not be extrapolated across vegetation layers or between composition and diversity patterns.

## Introduction

Although the effects of environmental or abiotic filtering on plant communities is often reflected in their species composition and richness, it is believed that this filtering mainly acts on the plants’ functional traits, rather than directly on the species’ identities (Lavorel and Garnier 2002, Kraft et al. 2015). Many studies have consequently observed strong trait – environment relationships across ecosystems (Wright et al. 2005, Ordoñez et al. 2009, Bruelheide et al. 2018). Not only functional trait composition, but also functional diversity can be affected by environmental filtering (Aros-Mualin et al. 2021). Functional diversity is, more specifically, expected to be reduced under environmentally stressful conditions, since only a limited number of functionally similar species will be able to establish (cf. trait underdispersion) (Weiher and Keddy 1995). The spatial extent at which environmental filtering occurs furthermore seems to differ among different environmental factors (Mokany and Roxburgh 2010, Bruelheide et al. 2018). While climatic factors mainly drive differences in species and trait composition across relatively large spatial scales, at smaller spatial scales, community (trait) composition is mainly structured by local-scale factors, such as variation in soil conditions (Bruelheide et al. 2018).

A recent study focusing on large-scale trait-environment patterns, suggested that the relative importance of different drivers even differs between woody and herbaceous species assemblages, with climatic variation more strongly impacting woody plant communities (Šímová et al. 2018). Consequently, within multi-layered forest ecosystems, environmental filtering might also differentially affect the woody overstory and herbaceous understory. For example, climate or (micro)climatic-related topography might more strongly impact the overstory, because the overstory is fully exposed to climatic variation, while the understory experiences buffered climatic variation under the protection of the forest canopy (Šímová et al. 2018, De Frenne et al. 2019). The species composition of the overstory might also be more likely to be filtered by more coarse-scale soil variation compared to that of the herbaceous understory, whose roots will be much more localized. The understory might additionally experience filtering due to small-scale environmental variation caused directly by variation in the overstory. Several studies have, for example, shown the impact of overstory related light availability and leaf litter on understory species composition (Komiyama et al. 2001, Wang et al. 2019, Majasalmi and Rautiainen 2020), trait composition (Maes et al. 2020) and functional diversity (Chabrerie et al. 2010).

Surprisingly little studies have, however, tried to quantify the similarities in environmental filtering between the over- and understory layers of forests (however see Ruokolainen et al. 2007, Rogers et al. 2008, Salazar et al. 2012). While this comparison is complicated for many temperate forest types, due to the often limited overstory species diversity, (sub)tropical forests offer an ideal study system to explore these relationships. In this study we focus on the proportional impact of several climate proxies (topography and ground fog frequency) and soil variation on the over- and the understory of the subtropical montane forests of northern Taiwan, along an elevation gradient ranging from 870 to 2130 m a.s.l.

Interestingly, the understory of subtropical montane forests contains a high diversity of fern and lycophyte species, next to angiosperms. For this reason, we focus specifically on the understory fern and lycophyte species in this study, while excluding angiosperms. Some studies have observed similar leaf trait-trait (Karst and Lechowicz 2007, Lin et al. 2020) and trait-environment relationships (Kessler et al. 2007, Kluge and Kessler 2007, Zhu et al. 2016, Campany et al. 2019) for ferns as for angiosperms, suggesting that functional patterns are similarly structured and thus comparable across both phylogenetic groups. Environmental filtering and trait-environment relationships nevertheless remain less well understood for fern and lycophyte communities, compared to angiosperm communities (Kessler et al. 2016). Effects of environmental filtering on understory fern community functional diversity has, for example, been observed in a few studies (Tanaka and Sato 2015, Zhang et al. 2017, Sessa et al. 2018), but not in others (Kluge and Kessler 2011, Aros-Mualin et al. 2021).

To allow optimal trait comparisons across both vegetation layers, we measured the same nine functional leaf traits for both overstory woody species and understory fern and lycophyte species. These nine traits were specifically chosen for their expected link to soil nutrient and/or climatic stress. Using this dataset, we addressed the following research questions:

– Do climate proxies and soil factors explain equal proportions of variation in the over- and understory community-level species and functional trait composition along the elevation gradient?
– Are these potential climate proxy and soil filtering processes also reflected in species and functional diversity along the elevation gradient?
– Can we find additional filtering of the understory species and trait composition due to variation in the overstory?

## Methods

### Study design

The study was performed in the Wulai district, New Taipei City, northern Taiwan, along an elevational transect ranging from Mt. Meilu (870 m a.s.l., 24.85°N 121.53°E) to Mt. Taman (2130 m a.s.l., 24.71°N 121.45°E, Fig. 1). The geological substrates mainly consist of argillite, shale, slate, sandstone and phyllite (Central Geological Survey, MOEA), with soils of low pH and high soil organic matter content. The study region is characterized by a humid subtropical climate (‘Cfa’ climate sensu the Köppen-Geiger system), with an average annual temperature of 16.1°C and average annual precipitation of 2070 mm (Lalashan weather station, 1374 m a.s.l., 24.68°N 121.40°E). Most precipitation falls during the summer, although the region is also affected by the north-eastern winter monsoon. The forest vegetation along the gradient varies from lower elevation *Pyrenaria-Machilus* subtropical winter monsoon forest, across mid-elevation *Quercus* montane evergreen broad-leaved cloud forest to higher elevation *Chamaecyparis* montane mixed cloud forest (Li et al. 2013).

**Figure 1.**
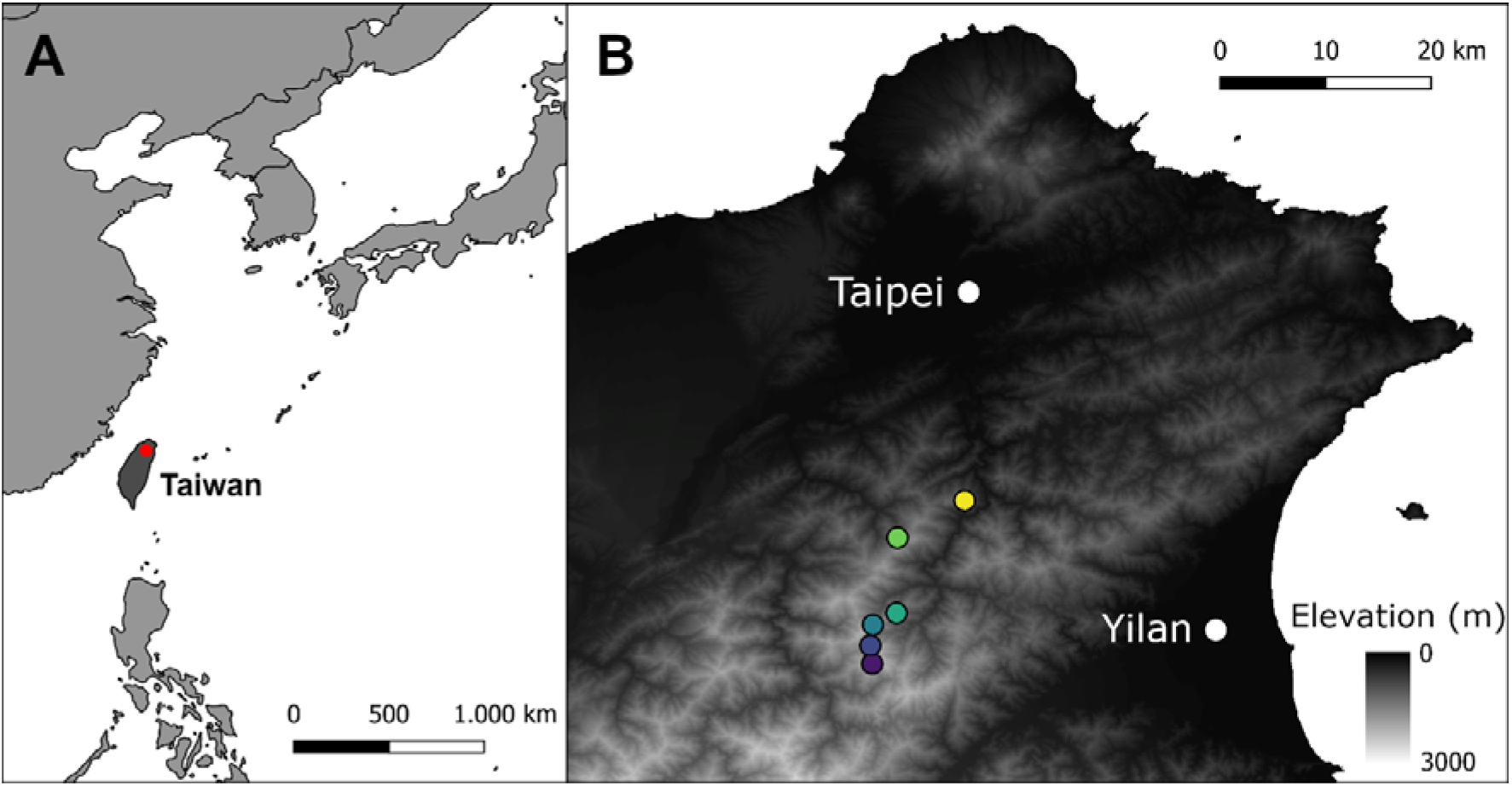
Location of the six elevation zones at which vegetation plots were established in northern Taiwan.

Along the transect, six elevation zones were delimited at 850, 1100, 1350, 1600, 1850 and 2100 m a.s.l. ± 50 m (Fig. 1). At each elevation zone, ten 10 m × 10 m plots were established across a secondary gradient in aspect and topography, ranging from the northeast facing (windward) to the southwest facing slope (leeward) across the ridge. Plots were positioned at least 50 m apart. Due to logistic constraints, only 9 plots were established at the 1850 m elevation zone, resulting in a total of 59 plots. For each plot, we recorded the presence of all woody species (angiosperm and gymnosperm shrubs and trees) taller than 2 m and with diameter in breast height (DBH) ≥ 1 cm (i.e. the ‘overstory’ vegetation). We also recorded the presence of all terrestrial (non-epiphytic) fern and lycophyte species, which together make up 66% of the plot-level herbaceous understory species richness in our study (i.e. the ‘understory’ vegetation).

### Climate proxies

We measured two topographic variables (i.e. elevation and heat load) that are known to relate to (micro)climatic conditions. The exact elevation of each plot was measured using GPS (GPSMAP 64st, Garmin, USA). To calculate heat load, we first measured the slope of each plot with a clinometer (SUUNTO PM-5/360 PC Clinometer, SUUNTO, Finland) from the upper plot edge, and the aspect of each plot with a compass (SILVA, Sweden). Aspect was then transformed into folded aspect, which was defined as the azimuth angle difference between the aspect and 45° (McCune and Keon 2002).We calculated heat load based on folded aspect and slope using equation 2 of McCune and Keon (2002). We additionally extracted average annual ground fog frequency for each plot from the ground fog frequency raster map for Taiwan (250 m per pixel resolution), developed by Schulz et al. (2017), based on MODIS satellite data. Ground fog frequency is expected to be an important environmental factor impacting cloud forest vegetation through effects on temperature, light availability, evapotranspiration and water availability (Fahey et al. 2016).

### Soil variables

For each plot, four soil samples of the top 0-10 cm were collected and pooled together for soil analysis. Each pooled soil sample was analyzed for pH, carbon:nitrogen ratio, total nitrogen, and phosphorous, potassium, magnesium, zinc, calcium, manganese, copper and iron content. See Appendix S1 for details of the soil chemical analysis.

To prevent collinearity among soil predictor variables, we performed a principal component analysis (PCA) on all measured soil chemical variables (after logarithmic transformation of Ca, C:N ratio, Mg, Mn, P and Zn and subsequent standardization to zero mean and unit standard deviation for all variables, Appendix S2). The three retained soil PC axes together explained 81.4% of the total variation. The first axis (termed ‘soil NPK’ in the text) reflected a gradient in nitrogen, phosphorous and potassium, next to several other micronutrients. The second axis (‘soil pH’) reflected an increase in pH and soil manganese, while the third axis (‘soil Cu’) was most strongly related to soil copper (positively) and calcium and iron (negatively) (Appendix S2).

We additionally measured soil depth at four positions in each plot using a 30 cm long soil depth meter (diameter 0.6 cm) and averaged values per plot. Soil more than 30 cm deep was recorded as 35 cm (16.9 % of the plots). Soil rockiness (the percentage content of rocks in the top 0-10 cm of the soil) was also estimated.

### Functional traits

Nine leaf traits were measured for 91 overstory woody species (sampled during 10/2014 and 12/2016-09/2018) and 48 understory fern and lycophyte species (sampled during 5/2017-10/2018), including all common species present in the plots and covering 74.0% and 63.2% of our over- and understory species pools, respectively (Appendix S3). The nine measured traits consisted of specific leaf area (SLA, mm^2^/mg), leaf dry matter content (LDMC, mg/g), area-based leaf chlorophyll content (SPAD units), leaf nitrogen content (leaf N, mg/g), leaf area (cm^2^), leaf thickness (Lth, mm), equivalent water thickness (EWT, mg/mm^2^), leaf ^13^C/^12^C stable isotope ratio (δ^13^C, ‰) and leaf ^15^N/^14^N stable isotope ratio (δ^15^N, ‰). Note that the first four traits are related to the leaf economics spectrum (LES) (cf. Wilson et al. 1999, Wright et al. 2004). Trait measurements largely followed standard protocols (Pérez-Harguindeguy et al. 2013), with some modifications for fern and lycophyte species. See Appendix S1 for trait measurement details.

These traits are expected to vary along climatic and soil variation gradients, because of their expected links to either soil nutrient stress (LES traits, δ^15^N, Lth; Wright and Cannon 2001, Hodgson et al. 2011), temperature or climatic stress (LES traits, leaf area; Wright et al. 2005, Dong et al. 2020) or drought (leaf area, Lth, EWT, δ^13^C; Medeiros et al. 2019, Maréchaux et al. 2020). While high values of δ^13^C are known to reflect high long-term water-use efficiency, and thus drought tolerance (Farquhar et al. 1982, Pérez-Harguindeguy et al. 2013), δ^15^N relates to a plant’s nitrogen acquisition strategy (Craine et al. 2015). More specifically, δ^15^N values around 0 ‰ usually indicate nitrogen fixation, while values around −2, −3 and −6 ‰ indicate plant nitrogen acquisition through arbuscular, ericoid and ectomycorrhiza, respectively (Craine et al. 2015). Note that EWT expresses the water mass content of a fresh leaf per unit leaf area, and is sometimes also called ‘succulence’ (Mantovani 1999, Féret et al. 2019).

All leaf level trait values were averaged at the species level after exclusion of leaf-level outliers (Z-score > 2.5 at the species level), as we assumed these values to most likely occur from measurement errors. This resulted in the exclusion of 0.61 % and 0.16 % of the trait values from the full leaf × trait matrix for the over- and understory, respectively. For leaf area we did not remove ‘outliers’, since all trait values could be verified for measurement errors. Missing trait values for nine overstory species were replaced by mean trait values across all overstory species, prior to further data analysis.

### Data analysis

We calculated the plot-level community mean (CM) values for each trait, as the average trait across all species present in the plot. Since only presence-absence data was collected, CM trait values were not weighted by species abundance. Species richness was calculated as the number of species present in the plot. We calculated two measures of functional diversity for each plot. Scheiner et al. (2017) has recently proposed to express functional diversity by the two parameters trait dispersion (M’) and trait ‘evenness’ or equability (^1^E(T)), next to species richness. While M’ quantifies ‘magnitude’, i.e. the amount of difference in trait values among species in a community, ^1^E(T) quantifies ‘variability’, i.e. the extent to which species are equally different from each other in trait values. Both measures are based on pairwise trait dissimilarities. Unlike the more traditionally used functional diversity parameters (i.e. functional richness, evenness and divergence), M’ and ^1^E(T) are independent from species richness and evenness, and thus solely reflect trait magnitude and variability, respectively (Scheiner 2019, Kosman et al. 2021). M’ and ^1^E(T) were calculated with the R script provided by Malavasi et al. (2018), based on the formulas of Scheiner et al. (2017), using Gower dissimilarity on the species × trait matrix (with traits standardized to Z-scores) to construct the trait dissimilarity matrix. Leaf area was logarithmically transformed prior to CM and functional diversity calculation. CM δ^15^N was additionally transformed using the equation (x + 4)^2^ for the overstory dataset and log (x + 2.5) for the understory dataset to obtain symmetrical distribution.

Before statistical analysis, we calculated variance inflation factors (VIF) to identify potential collinearity issues, separately among the different climate proxies and among soil variables. Collinearity was identified for folded aspect and heat load (VIF > 5), and thus we excluded folded aspect from all statistical models. We consequently retained three climate proxies (elevation, heat load and ground fog frequency) and five soil variables (soil depth, soil rockiness and three soil PC axes, namely soil NPK, soil pH and soil Cu). Heat load was squared and soil rockiness was square root transformed prior to statistical analyses to improve symmetrical distribution of their values.

To test the effects of climate proxies and soil variables on the over- and understory species composition, we performed separate redundancy analyses (RDA) on the respective plot × species matrices. These RDA models were repeated on the plot × species matrices including only species for which traits were measured. Similar RDA models were performed for the over- and understory plot × CM trait matrices, with CM of traits standardized to Z-scores (CWM-RDA, Nygaard and Ejrnæs 2004). The overstory trait RDA model was also repeated after excluding all deciduous species, to assess the potential effect of deciduous species on the trait patterns.

After assuring global significance of each of the global RDA models containing all environmental predictors, we performed variation partitioning on each model to assess the proportions of variation in each plot × species or plot × CM trait matrix explained by either climate proxies or soil variables, expressed by adjusted R^2^. Next, we performed forward model selection among all environmental predictors, based on adjusted R^2^-values (conditional effects) and their significance assessed using Monte Carlo permutation tests (9999 permutations). This model selection allowed us to assess which environmental variables were most important predictors for each species and CM trait dataset. We are aware that the CWM-RDA method is prone to inflated Type I error rate when using the standard Monte Carlo permutation test (Šmilauer and Lepš 2014, Zelený 2018). Since there is no published solution to this problem, however, we nonetheless use these tests. Before forward model selection, we again used VIFs to ensure that no collinearity occurred among any of the combined climate and soil variables. All multivariate analyses were performed with the ‘vegan’ R package (Oksanen et al. 2017).

We additionally performed partial redundancy analysis for the understory plot × species and plot × CM trait datasets, to assess the potential additional effects of the overstory. To achieve these models, we first performed two PCA’s, one on the overstory plot × species matrix and one on the overstory plot × CM trait matrix. For both models, we only retained PCA axes for which the eigenvalue was larger than the average inertia (Kaiser-Guttman criterion, Ibanez 1973). Hence, we retained 15 and 3 axes for the overstory species and CM trait PCA’s, respectively. Next, we performed a partial RDA (pRDA) on the understory plot × species matrix with all retained overstory species PCA axes as explanatory variables, while partialling out the effects of all measured environmental variables, and assessed the model’s overall significance. Similarly, we assessed the effect (including overall significance) of the retained overstory trait PCA axes on the understory plot × trait matrix while partialling out all climate proxies and soil variables.

The effects of climate proxies and soil variables on species and functional diversity were assessed using generalized linear models. For species richness (a count variable) we used a Poisson probability distribution and log link function. For M’ and ^1^E(T) we used a gamma probability distribution with inverse link function, since these variables consist of positive, continuous, right skewed data. Due to the nonlinear relationship between elevation and understory species richness, and fog frequency and overstory ^1^E(T), we included the quadratic term for elevation and fog frequency to the respective models. We performed variation partitioning on the full diversity models to quantify the proportion of explained deviance by climate proxies and soil variables. Each full model was then reduced using a AIC-based comparison of all predictor-subset combinations of the full model, using the ‘dredge’ function in the ‘MuMIn’ R package (Barton 2019). Model assumptions were checked using the ‘DHARMa’ R package (Hartig 2021).

If traits and diversity are shaped by similar environmental drivers for the under- and overstory along our gradient, we expect them to be positively correlated between vegetation layers. To verify this, we performed simple linear regressions between each understory CM trait (response) and overstory CM trait (predictor). Based on the scatterplots for these regressions, we also included a quadratic term for the CM SLA model (i.e. understory CM SLA ~ overstory CM SLA + (overstory CM SLA)^2^). These regressions were also performed for overstory CM traits excluding deciduous species. Similar regressions were additionally constructed for S, M’ and ^1^E(T) between under- and overstory. All p-values of the pairwise regressions were corrected for type-I error inflation using the false discovery rate method (Benjamini and Hochberg 1995) with the ‘p.adjust’ function in the ‘stats’ R package. All analyses were performed with R version 4.0.5.

### Results

Species composition of both over- and understory species was significantly affected by climate proxy and soil variation, together explaining 29.9% and 30.7% of the total variation (i.e. adjusted R^2^), respectively (Fig. 2A&B). Variation partitioning indicated that for both over- and understory species composition, climate proxies explained a higher relative proportion of this variation (overstory: 80.4%, understory: 85.0%) than soil (overstory: 54.9%, understory: 54.2%). However, for both vegetation layers, around one third of the explained variation was shared by climate proxies and soil (Fig. 2A&B). The final RDA models after forward model selection retained all climate proxy variables and soil pH for both the overstory and understory species composition. For the overstory species composition, soil NPK and soil Cu were additionally retained (Table 1, Fig. 3A&B). RDA and variation partitioning results were barely affected when RDA was performed on the plot × species matrices including only species for which traits were measured (Appendices S4 & S5). The pRDA indicated that an additional 7.1% (p < 0.001) of the total variation in understory species composition was explained purely by overstory species composition.

**Table 1.**
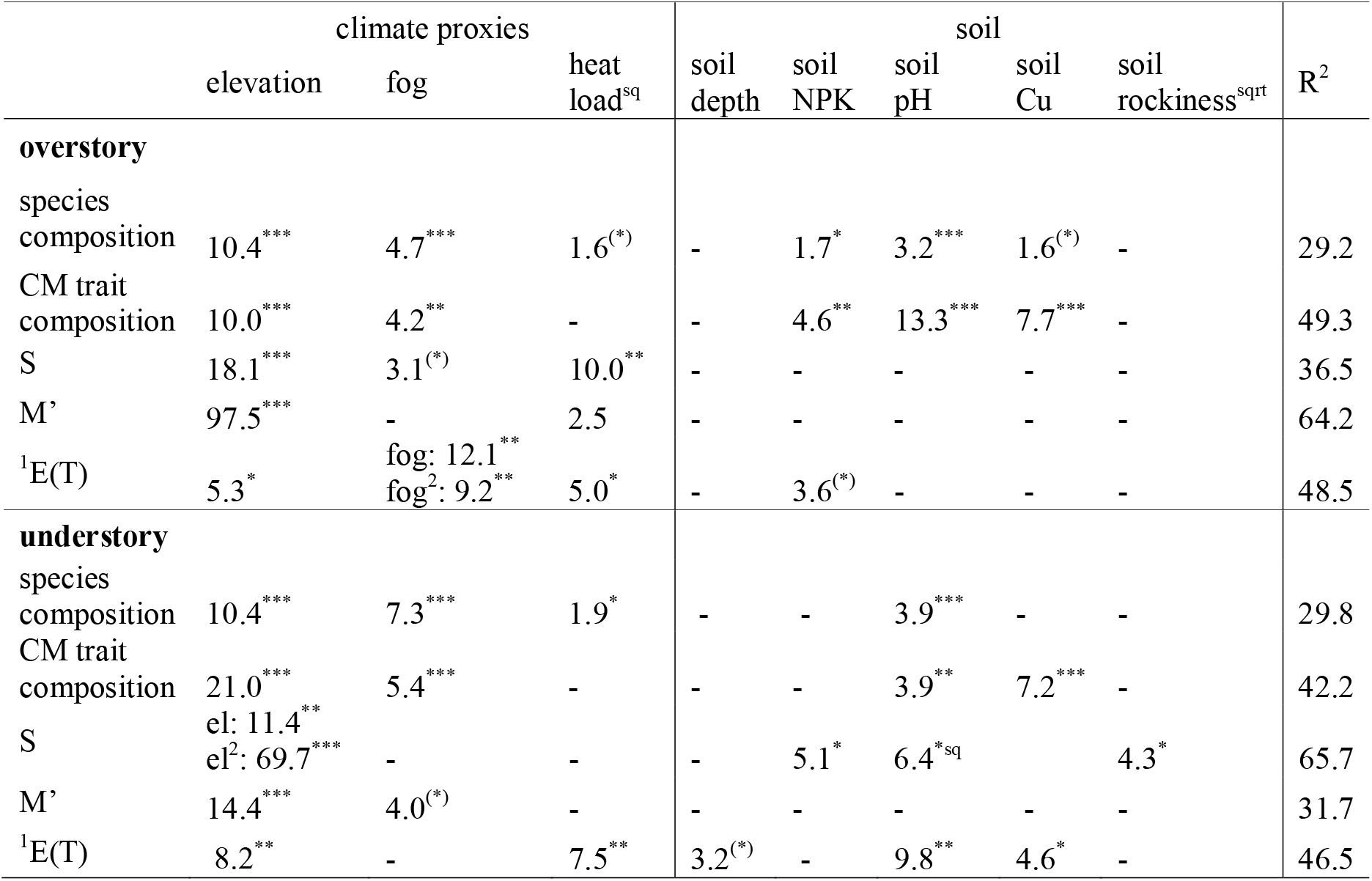
Parameter estimates of the reduced redundancy analyses (RDA) for species and trait composition and the reduced generalized linear models for species richness (S), functional divergence (M’) and functional equability (^1^E(T)), for the overstory and understory datasets separately. Test statistic (F) for each retained predictor and full model adjusted R^2^ provided. For **s**oil principal components ‘soil NPK’, ‘soil pH’ and ‘soil Cu’, see Table 1. ^(*)^0.10 ≥ p-value > 0.05; ^*^0.05 ≥ p-value > 0.01; ^**^0.01 ≥ p-value > 0.001; ^***^0.001 ≥ p-value. ^sqrt^ = square root transformation, ^sq^ = squared transformation of environmental factor. El = elevation, fog = ground fog frequency.

**Figure 2.**
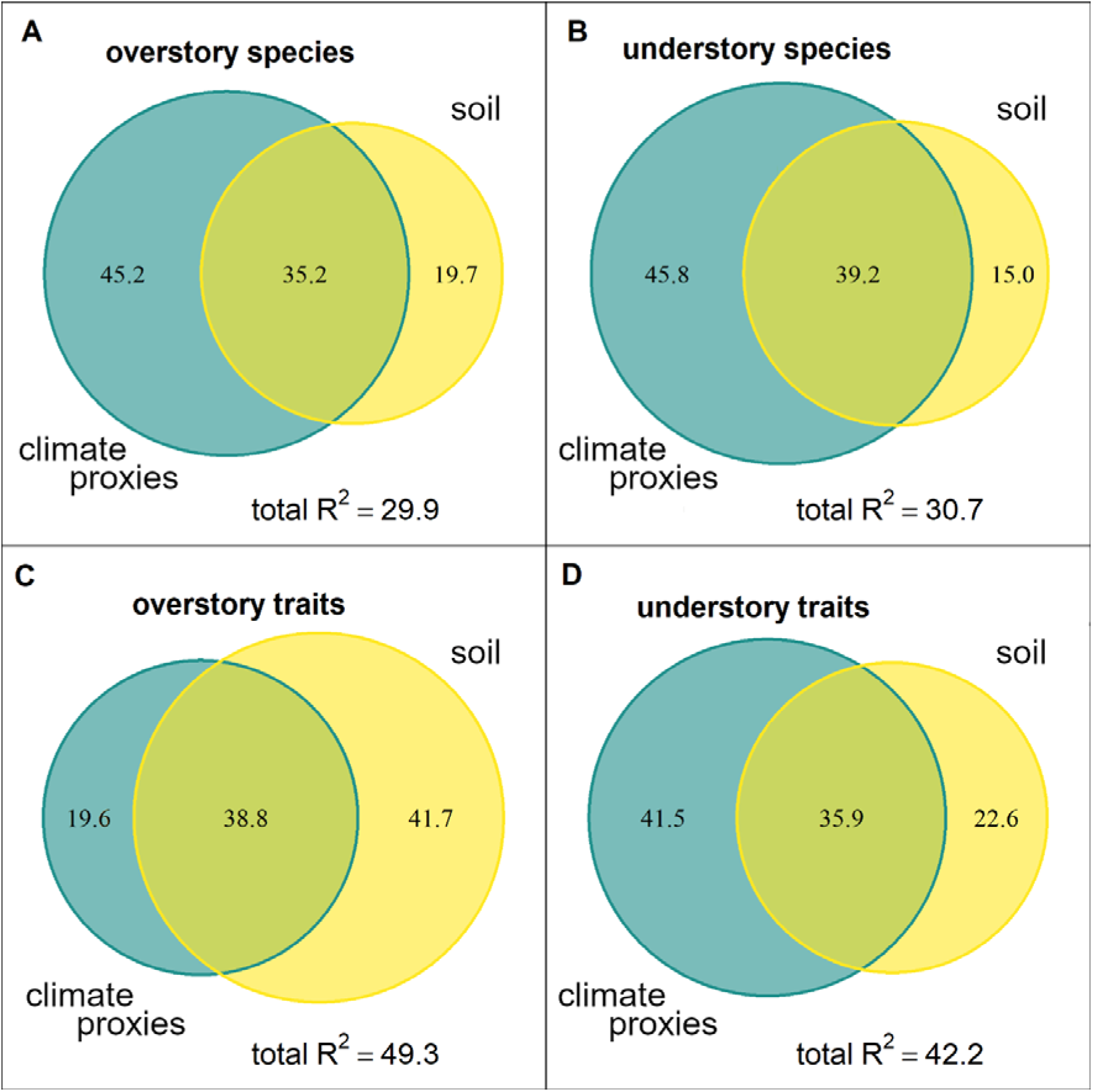
Venn diagrams visualizing the variation partitioning between climate proxy and soil variable effects on A. the overstory plot × species matrix, B. the understory plot × species matrix, C. the overstory plot × community mean (CM) trait matrix, D. the understory plot × CM trait matrix, using redundancy analysis (RDA). Numbers in the Venn diagrams correspond to the relative proportions of the total explained variation. The total explained variance (adjusted R^2^) is also presented.

**Figure 3.**
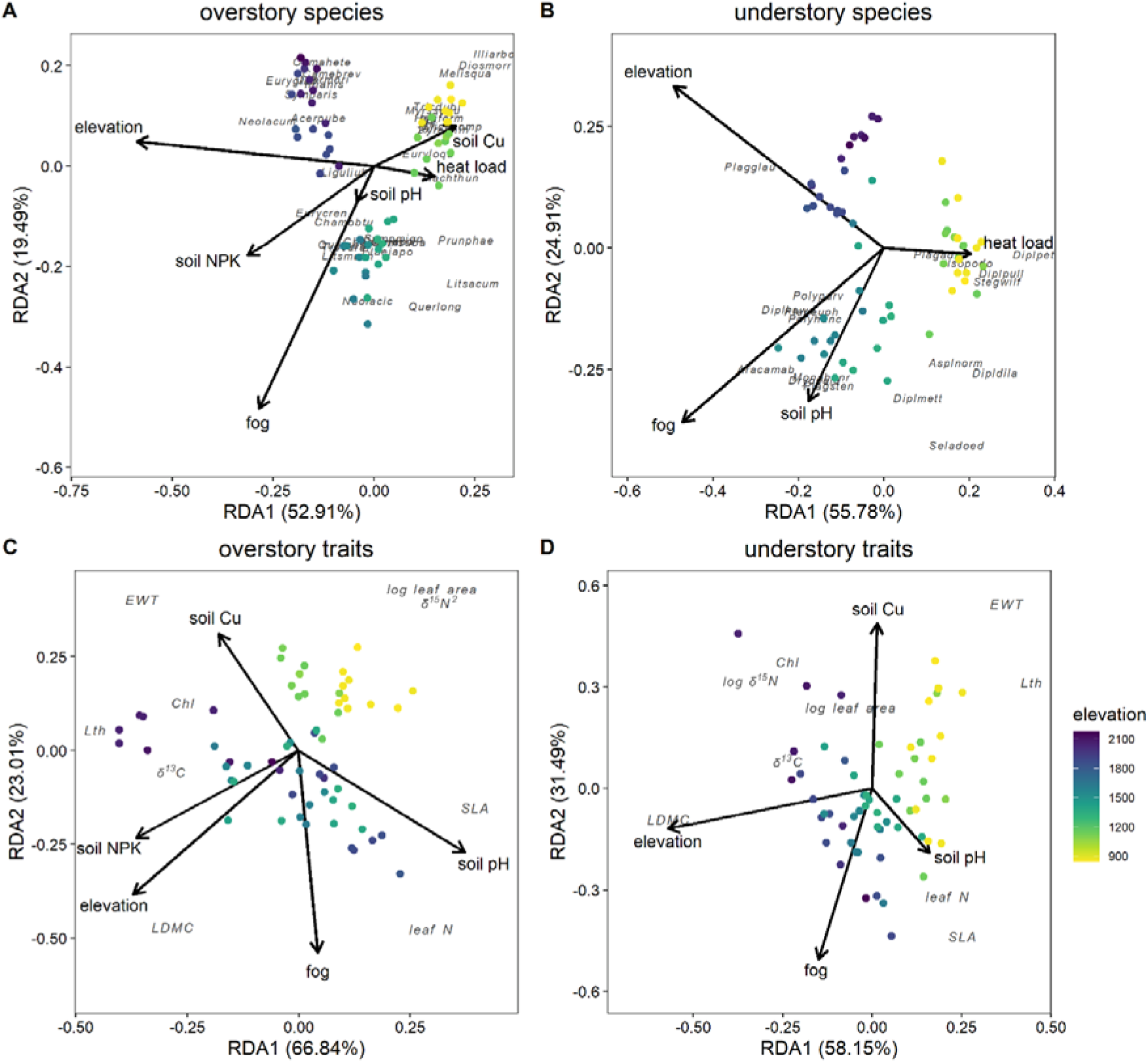
Triplots for the redundancy analyses (RDA) after forward model selection, exploring the effect of environment (climate proxies and soil) on A. the overstory plot × species matrix, B. the understory plot × species matrix, C. the overstory plot × community mean (CM) trait matrix, D. the understory plot × CM trait matrix. Plots visualized as points, with colors indicating plot elevation, 40% most common species with 20% best fit to the RDA axes are visualized as codes (see Appendix S3), CM trait vectors visualized as vector tips with names in italics, environmental variables visualized as vectors. Note that ‘soil NPK’, ‘soil pH’ and ‘soil Cu’ refer to the first three soil PCA axes, respectively (Table 1). Chl = leaf chlorophyll content, δ^13^C = the leaf ^13^C/^12^C stable isotope ratio, δ^15^N = the leaf ^15^N/^14^N stable isotope ratio, EWT = equivalent water thickness, Lth = leaf thickness.

CM trait composition variation was also related to climate proxies and soil for both over- (adjusted R^2^ = 49.3%) and understory species (adjusted R^2^ = 42.2%) (Fig. 2C&D). Variation partitioning showed that climate proxies were the most important predictors of CM trait variation for understory species (77.4% of the total explained variation for climate proxies vs. 58.5% for soil), while soil explained most variation for overstory CM traits (58.4% for climate proxies vs. 80.5% for soil). For both vegetation layers’ trait composition, around one third of the explained variation was shared by climate proxies and soil (Fig. 2C&D). After forward model selection, elevation, ground fog frequency, soil pH and soil Cu were retained for both CM trait RDAs, while soil NPK was additionally retained for the overstory CM trait RDA (Fig. 3C&D). The RDA results were largely similar for overstory traits excluding deciduous species. However, climate proxies became more important than soil after variance partitioning (Appendices S4 & S5).

The results of the CM trait RDAs furthermore suggest an increase in CM LDMC with elevation for both vegetation layers and a decrease in CM of leaf area and δ^15^N with elevation for the overstory. High ground fog frequency seems to result in lower CM EWT for both vegetation layers, while high soil pH seems to result in high CM of SLA and leaf N and low CM Lth for the overstory, and high CM leaf N for the understory (Fig. 3C&D). Note that these patterns can be inferred from Fig. 3 because the involved environmental variables and traits were well represented by the first two RDA axes (high axis loadings). Evaluation of the third RDA axis loadings for the understory furthermore suggests that high soil pH is also related to high CM leaf N for the understory (results not shown). The pRDA showed that the overstory trait composition could explain an additional 4.5% (p = 0.011) of the variation in the understory trait composition after partialling out the effects of climate proxies and soil variables.

CM of LDMC and leaf N were positively related, while CM of Lth and δ^15^N were negatively related between both vegetation layers. CM SLA showed a parabolic relationship, with the highest values for the understory at intermediate values for the overstory species. CM of leaf area, chlorophyll content, δ^13^C and EWT, on the other hand, were not significantly related between both vegetation layers (Fig. 4, Appendix S6). Excluding deciduous species for the overstory weakened, but did not change the direction of all trait - trait relationships, except for leaf thickness, for which the negative relationship strengthened (Appendices S6 & S7).

**Figure 4.**
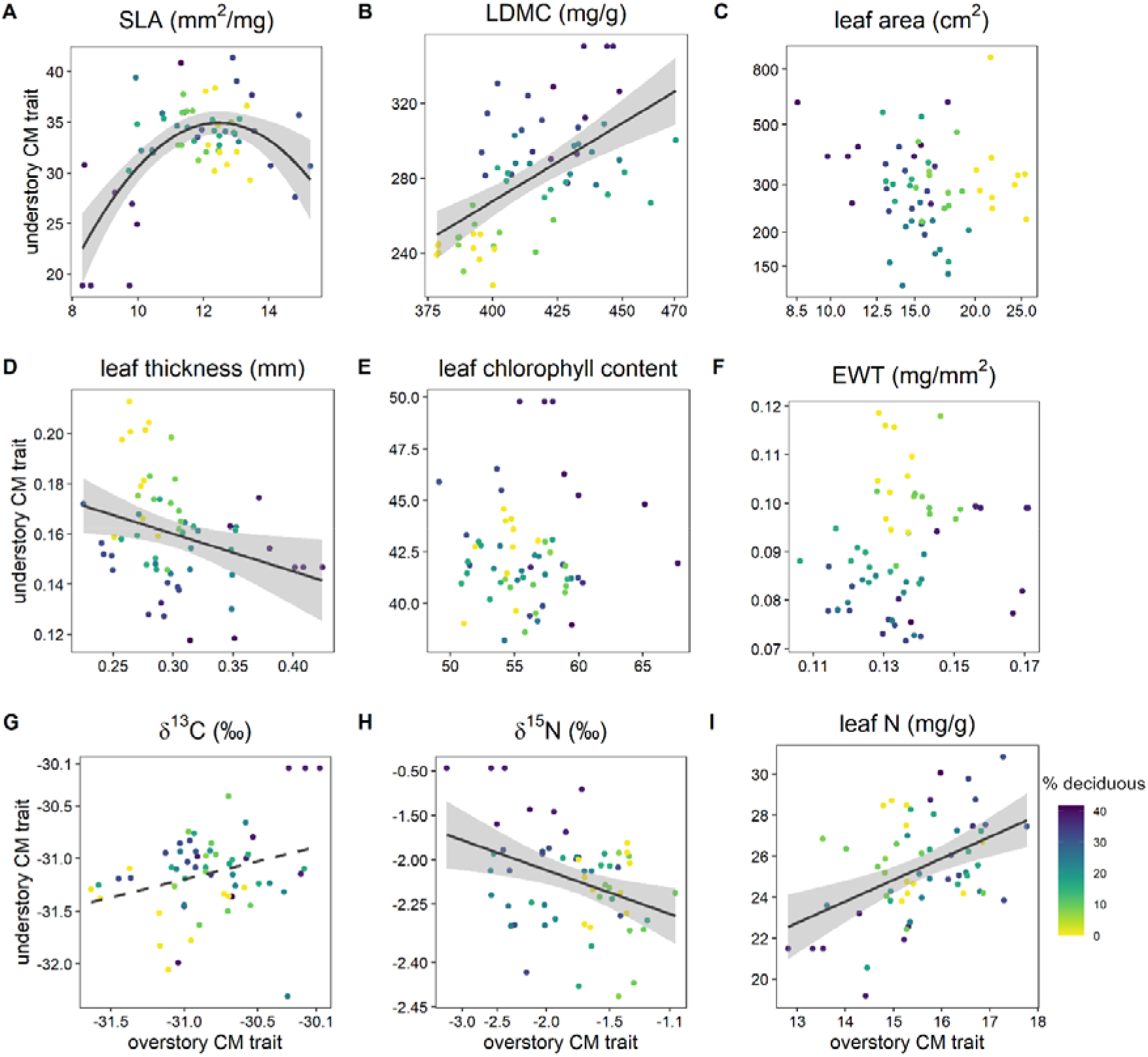
Scatterplots for pairwise regressions between plot-level overstory and understory community mean (CM) trait values. Solid regression line + SE presented for significant regressions, dashed line for marginally significant regression (see Appendix S6). Each datapoint corresponds to one vegetation plot, with colors indicating plot elevation. δ^13^C = the leaf ^13^C/^12^C stable isotope ratio, δ^15^N = the leaf ^15^N/^14^N stable isotope ratio, EWT = equivalent water thickness, Lth = leaf thickness.

Compared to soil, climate proxy variables explained more of the variation in species richness, functional divergence and functional equability for both vegetation layers (Table 1, Appendix S8). Soil variation nonetheless contributed a small additional proportion to the total explained variation (< 20%) in the understory species richness and functional equability for both vegetation layers (Table 1, Appendix S8). While elevation was the strongest climate proxy predictor for all diversity measures of both the over- and understory, the direction of these relationship differed between both vegetation layers. While overstory species richness declined with elevation, understory species richness was highest at intermediate elevation (Fig. 5A). Functional dispersion increased with elevation for both vegetation layers (Fig. 5C), while functional equability increased with elevation for the overstory, and decreased for the understory (Fig. 5E). Soil pH was the strongest soil predictor for understory species richness (positive relationship) (Fig. 5B) and functional equability (negative relationship) (Fig. 5F).

**Figure 5.**
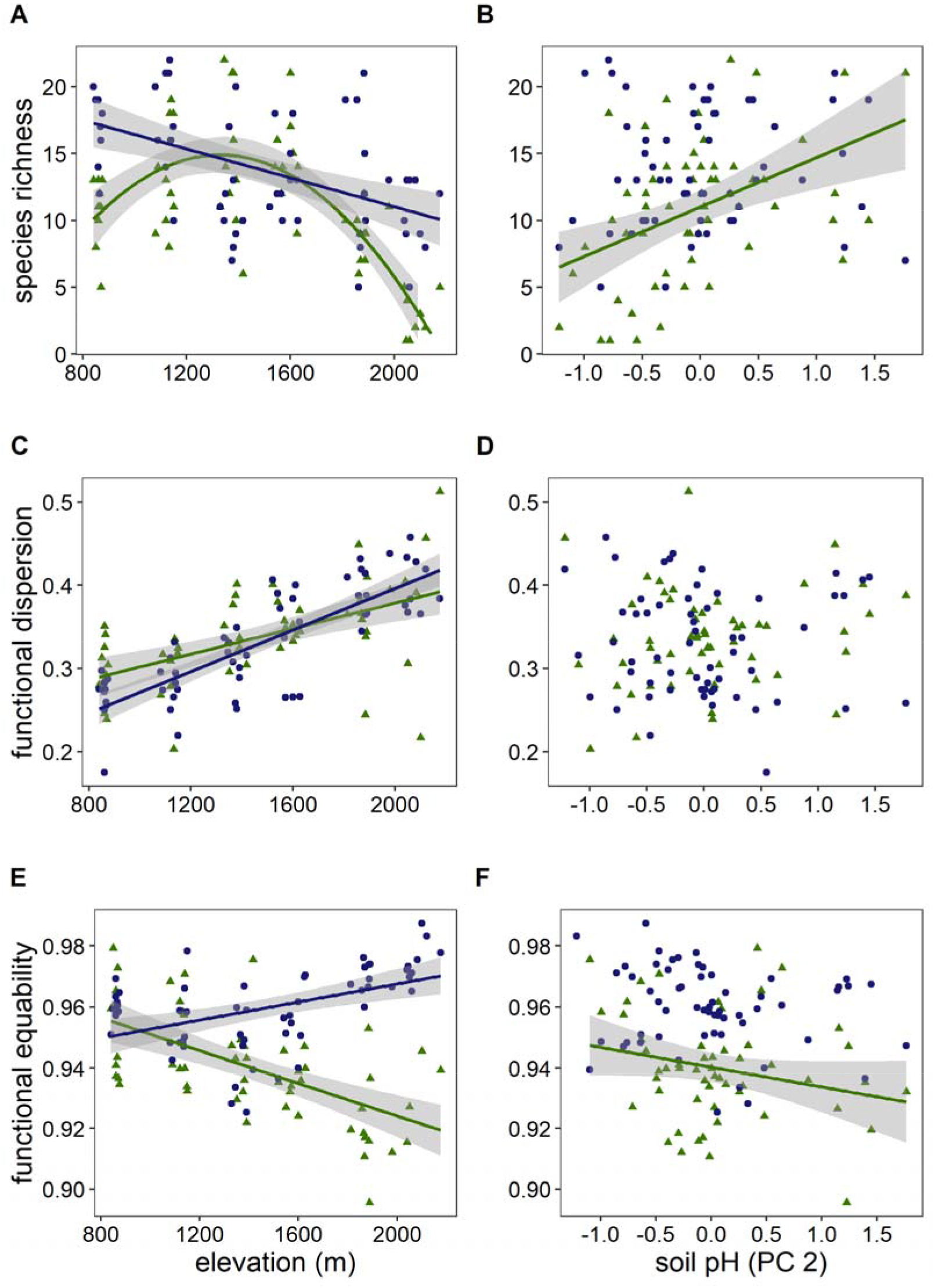
Scatterplots between three species and functional diversity measures, on the one hand and elevation and the second soil PCA axis (soil pH), on the other hand. Blue circles and lines = overstory species, green triangles and lines = understory species. Solid regression line + SE presented for significant regressions (Table 1).

Species richness and functional equability were not significantly related for both vegetation layers, while functional dispersion was significantly positively related between the over- and understory (Appendices S6 & S9).

## Discussion

### Species and trait composition

Climate proxies were more strongly related than soil to the species composition of the overstory. This result was expected, since, unlike the understory, the overstory is fully exposed to climatic variation (Šímová et al. 2018, De Frenne et al. 2019). Surprisingly, however, also the species and trait composition of the fern and lycophyte understory were mainly shaped by climate proxies. This nonetheless agrees with the literature, where climate often strongly impacts fern trait composition (e.g. Kessler et al. 2007, Kluge and Kessler 2007, Sessa and Givnish 2014). It has been suggested that the limited potential for controlling evaporation makes ferns more sensitive to drought than angiosperms (Brodribb and Holbrook 2004, Zhang et al. 2014). This could thus explain why elevation (temperature) and ground fog frequency (relative humidity) rather than soil variation were the main drivers of fern species and trait composition in our study.

The high relative importance of elevation is not surprising, considering the quite steep elevational gradient of 1260 m in our dataset. The trait responses followed expectations of more resource conservative (e.g. high LMDC) and smaller leaves with increasing elevation for both vegetation layers (Wright et al. 2005, Dong et al. 2020). The high ground fog frequency likely increased water availability, as suggested by drought-related trait states (high EWT for both vegetation layers and Lth for understory) in low fog plots in our study (cf. Medeiros et al. 2019, Maréchaux et al. 2020). Frequent fog can, however, additionally reduce light availability (up to 10-50%) and local temperature (up to 3-6°C) (Lai et al. 2006, Reinhardt and Smith 2008) and negatively impact photosynthesis by preventing leaf transpiration and promoting the growth of epiphyllous lichens and algae. While these conditions are highly suitable for fern species, which often seem adapted to low light and high water availability (Sessa and Givnish 2014, Hernández-Rojas et al. 2020), they likely present less suitable growing conditions for most tree species by hampering photosynthesis (Fahey et al. 2016). These potential different responses of both vegetation layers were, however, not reflected in their respective trait composition. More detailed future work using field-based fog or relative humidity measurements should be used to further explore these potential effects.

Soil did nevertheless still impact the species and trait composition of both vegetation layers and was, unexpectedly, more important than climate proxies in structuring overstory trait composition. The higher importance of soil for overstory traits compared to species composition was not due to differences in the species included in both models, since the species RDA model including only species for which traits were measured gave similar results. The ‘soil pH’ ordination axis was the most important soil driver, and likely better reflects plant nutrient availability than soil NPK in the typically highly acidic soils with low decomposition rates and high soil organic matter of cloud forests (Fahey et al. 2016). The high nutrient levels following standard soil analysis of these soils likely reflect nutrients trapped in undecomposed organic matter, rather than plant available nutrients. This could explain the presence of acquisitive leaf traits (high SLA and leaf N) in less acidic soils for both vegetation layers (Wright and Cannon 2001). The lower relative importance of soil variables compared to climate proxies for the understory, on the other hand, might be because understory fern species respond to more small-scale soil variation than that measured at the plot-level in this study.

Interestingly, if deciduous species are excluded from the overstory, climate proxies become the most important driver of overstory trait composition, due to reduced importance of soil pH. Deciduous woody species are more common on steep wind-exposed slopes in northern Taiwan, probably because of their ability to avoid environmental stress during winter by shedding their leaves. These steep slopes usually have higher soil pH in comparison with less steep of flat ridges at the same elevation, perhaps due to surface erosion removing more acidic soil and litter and increasing availability of more cation-rich weathered parental rock material. This contradiction between seemingly stress-adapted niches and more acquisitive LES traits (cf. low leaf longevity, Wright et al. 2004) of deciduous species, makes it difficult to assess if these soil pH patterns are caused by LES nutrient-availability or wind-exposure and slope. This shows how soil, topography and climate can interact in their impact on species- and trait composition, as also illustrated by the high overlap in explained variation by climate proxies and soil for all variation partitioning analyses.

We found that the overstory can act as an additional filter on the understory’s species and trait composition, through potential light availability or leaf litter effects, as observed in previous studies (Komiyama et al. 2001, Wang et al. 2019, Maes et al. 2020, Majasalmi and Rautiainen 2020). The observational nature of our study does, however, not allow us to assess the causality of this effect. Alternatively, the relationship between overstory and understory could be caused by unmeasured environmental conditions affecting species and trait composition of both layers simultaneously.

Despite some similarities in the environmental responses of both vegetation layers, only two of the nine measured traits showed a significant positive relationship between over- and understory. This illustrates that environmental filtering differs substantially for trees and ferns and is most clearly illustrated by Lth. For ferns, Lth seemed to respond to drought (cf. Kluge and Kessler 2007), but was linked to low nutrient availability in trees (cf. Read et al. 2006), resulting in a negative correlation between both. Direct impact of the overstory on the understory could also have shaped unexpected trait relationships. The negative correlation for δ^15^N might, for example, reflect niche differentiation among vegetation layers to prevent competition for different nitrogen sources. The quadratic relationship for SLA could also be due to overstory impact on the understory. While nutrient availability mainly structured SLA for the overstory, for the understory, SLA might be affected by a combination of nutrient limitation under low overstory SLA (cf. Kessler et al. 2007) and more strong light competition due to shading under high overstory SLA.

### Species and functional diversity

Species richness was most strongly affected by elevation for both vegetation layers. Both the decrease in overstory species richness and humped-shaped relationship for understory fern species along elevation are consistent with previous studies in (sub)tropical forests (Kluge and Kessler 2011, Qian and Ricklefs 2016, Hernández-Rojas et al. 2020). At lower elevation, lower water availability is expected to reduce diversity of drought-sensitive lifeforms such as ferns and lycophytes (Kessler et al. 2011, Weigand et al. 2020). At higher elevations, on the other hand, species richness of both ferns and angiosperms will be reduced by the stronger climatic stress associated with lower temperatures (Kessler et al. 2011). Interestingly, soil explained a much higher proportion of variation in understory than overstory species richness, thus showing the opposite pattern as observed for species and trait composition. The impact of soil productivity (soil pH), next to climate on fern species richness is nonetheless in agreement with previous work (Tuomisto et al. 2014, Weigand et al. 2020). Not surprisingly, species richness was not correlated between the two vegetation layers, further illustrating that species richness is shaped by different environmental drivers for overstory trees and understory ferns and lycophytes.

Functional diversity patterns suggested that for the overstory, the community trait composition is not experiencing increased trait convergence among species with elevation, as expected under increased environmental filtering (Weiher and Keddy 1995). On the contrary, species trait overlap seems to be reduced, resulting in higher average trait distances among species (M’), combined with high equal spacing of species in the trait space (^1^E(T)), a pattern usually attributed to increased importance of competition among species (Kraft et al. 2008). Functional equability was nonetheless highest at intermediate ground fog frequency levels, potentially reflecting trait clustering (environmental filtering) due to drought stress and high air humidity at each respective end of the ground fog frequency gradient.

For the understory, trait composition seemingly clustered in separate distinct trait sets, potentially reflecting (environmental) filtering of a few alternative trait combinations for ferns at high elevation (increasing M’, decreasing ^1^E(T)). A similar pattern was also observed for terrestrial fern communities along four tropical elevation gradients (Aros-Mualin et al. 2021). Elevation related most strongly to functional dispersion in both vegetation layers and functional equability for the overstory. This again mirrors the results of Aros-Mualin et al. (2021), who found that temperature was the most important predictor of fern functional diversity. Functional equability of the understory, on the other hand, was almost equally strongly affected by soil variation and climate proxies. This soil (pH)-driven environmental trait filtering for the fern understory (cf. Zhang et al. 2017, Sessa et al. 2018), but not the overstory, thus mirrors the impact of soil on the species richness patterns in our study.

## Conclusions

Both climate proxies and soil were important predictors of species and trait composition of both vegetation layers. The stronger effects of climate proxies for understory ferns and lycophytes compared to overstory trees is likely due to their higher vulnerability to drought. The environmental drivers furthermore seem to affect very different response traits in both vegetation layers, which together with additional overstory effects on understory traits, results in a disconnection of community-level trait values across layers. Interestingly, the relative importance of soil and climate proxies on species or trait composition cannot be extrapolated to species or trait diversity, which showed very different patterns. This study illustrates that environmental filtering can differentially affect species, trait and diversity patterns and can be highly divergent for forest overstory and understory vegetation.

## Acknowledgements

We would like to thank to all volunteers who contributed to the field work and lab trait measurements. Hsin-Yen Teng and Kun-Sung Wu helped with the logistics of the field work, specimen determination and dataset preparation. This study was supported by Ministry of Science and Technology, Taiwan (106-2621-B-002-003-MY3 and 109-2811-B-002-644).

