## Appendix for "Do forest over- and understory respond to the same environmental variables when viewed at the taxonomic and trait level?"

### Appendices

#### Appendix S1. Extended methods section.

##### Soil chemical analysis

Soil samples were air dried and then sieved using a 2.0 mm sieve (Endecotts Ltd, UK) prior to further analysis. Soil pH was measured in a 1:1 soil-water solution (20g of soil was mixed with 20 ml of distilled water and occasionally stirred) using pH meter (Metrohm, 827 pH lab, Switzerland). The mixture was left to rest for 1h before pH measurement (Thomas, 1996). Soil extractable Ca, Cu, Fe, K, Mg, Mn, P and Zn content (mg/kg) were extracted by Mehlich No. 3 solution, and the concentration of the elements was assessed using inductively coupled plasma atomic emission spectroscopy (ICP-AES, JY2000, HORIBA Jobin Yvon, USA). More specifically, we mixed 5 g of soil with 50 ml of Mehlich No. 3 extracting solution (M3, Mehlich 1984). The solution was shaken for 5 min at 160 rpm and subsequently filtered twice using a Whatman No.42 filter paper and a filter membrane (<0.45µm). The resulting filtrate was used for ICP-AES analysis. Total soil carbon and nitrogen content (%) was assessed using the dry combustion method (SSLS, 2014). An air-dried (<180µm) sample was packed in a tin foil, weighed, and analyzed by an elemental analyzer (vario EL III Element Analyzer, Elementar Analysensysteme GmbH, Germany). Soil total carbon and total nitrogen content were subsequently corrected by taking into account the soil moisture content of the analyzed samples. Soil moisture content was assessed as the weight loss after over drying at 105°C for 24h. Soil C:N ratio was calculated by dividing total soil carbon by total soil nitrogen.

##### Trait measurements

For each woody species, we selected 3-5 individual plants, from which we collected a branch with several mature, healthy leaves each, and used three of them for trait measurement. Where possible, branches were obtained from the sun-exposed part of the canopy, using an 11 m long telescopic knife. For each fern and lycophyte species, we selected, on average, 5 (range 1-12) individual plants, from which we collected 1-6 fully expanded leaves (fronds) each. Only healthy, fully expanded fronds were collected, mainly from mature plants (i.e. plants containing sporophylls). For species with frond dimorphism, traits were measured on the sterile frond (trophophyll). For non-dimorphic species, traits were measured on sori-containing fronds. Most of the leaf samples (for both woody species and ferns) were collected in the study region, either in a vegetation plot or within a 50 m radius from a plot, and stored in wet sealed Ziploc bags at low temperature (< 10°C) until trait measurement. For species occurring across multiple elevation zones in our study, individuals were collected across its full range, to obtain representative trait measurements.

Leaf fresh weight (g) and area ($cm^2$) were measured on fresh, fully rehydrated leaves using a 0.1 mg precision balance (OHAUS Adventurer AR2140, USA) and scanner (Perfection V370 Photo, EPSON, Japan), respectively. Leaf thickness (Lth, mm) was measured on each rehydrated leaf using a digital thickness gauge with a 0.001 mm precision (DML digital thickness gauge, UK). After fresh weight, leaf area and Lth measurements, leaves were dried for at least 72h at 70°C. After drying, leaf dry weights (mg) were measured with the same balance. Specific leaf area (SLA, $mm^2/mg$) was then calculated by dividing the one-sided leaf area by the leaf dry weight and leaf dry matter content (LDMC, mg/g) as the leaf dry weight divided by the leaf fresh weight. Equivalent water thickness (EWT, $mg/mm^2$), which quantifies area-based water content, was calculated as (leaf fresh weight – leaf dry weight)/ leaf area (Mantovani 1999, Féret et al. 2019).

For each collected leaf we performed six random area-based chlorophyll measurements, three left and three right of the midvein. For all understory ferns and most overstory woody species, chlorophyll content was estimated using a SPAD-502 chlorophyll meter (KONICA MINOLTA, Japan). For a few overstory species chlorophyll measurements were performed with a CCM-200 chlorophyll meter (APOGEE, USA), which were subsequently converted to SPAD units based on a empirically defined calibration curve (SPAD = -12.85 + ln(CCM) × 18.26). The calibration curve was constructed by estimating chlorophyll content with both SPAD and CCM for 39 leaves of 13 overstory species.

For each species, one to four of the collected leaves were used to quantify leaf nitrogen content (leaf N, mg/g), leaf $^{13}C/^{12}C$ stable isotope ratio ($\delta^{13}C$, ‰) and leaf $^{15}N/^{14}N$ stable isotope ratio ($\delta^{15}N$, ‰). Leaf N was measured on grounded dry leaf material with a FlashEA 1112 series elemental analyzer (Thermo Fisher Scientific, Italy), while $\delta^{13}C$ and $\delta^{15}N$ were measured on grounded dry leaf material with a Delta V Advantage isotope ratio mass spectrometer (Finnigan Mat, Germany), using Peedee belemnite (PDB) and atmospheric nitrogen as global standards for $\delta^{13}C$ and $\delta^{15}N$, respectively.

**Appendix S2.** Soil variable axis loadings and cumulative soil variation represented by the three selected principal component axes (PC). Main contributors to each ordination axis in bold. All soil nutrients present soil extractable values. $^{l}$ = log transformed.

| | PC1 (soil NPK) | PC2 (soil pH) | PC3 (soil Cu) |
|---|---|---|---|
| C:N ratio$^{l}$ | **0.31** | -0.33 | 0.29 |
| Ca$^{l}$ | 0.28 | 0.26 | **-0.38** |
| Cu | 0.18 | 0.28 | **0.59** |
| Fe | -0.12 | -0.40 | **-0.51** |
| K | **0.36** | 0.02 | 0.02 |
| Mg$^{l}$ | **0.40** | -0.05 | 0.02 |
| Mn$^{l}$ | 0.12 | **0.59** | -0.30 |
| pH | -0.26 | **0.45** | 0.08 |
| P$^{l}$ | **0.38** | 0.06 | -0.14 |
| total N | **0.33** | 0.07 | -0.23 |
| Zn$^{l}$ | **0.39** | -0.17 | 0.05 |
| Cumul. variation (%) | 52.1 | 70.3 | 81.3 |

**Appendix S3. Full species list separated in over- and understory (fern and lycophyte) species**. The column 'traits' indicates for which species functional traits were measured.

| Nr | Species | Traits | Nr | Species | Traits |
|---|---|---|---|---|---|
| **Overstory species** | | | | | |
| 1 | *Acer kawakamii* | × | 35 | *Ficus formosana* | |
| 2 | *Acer morrisonense* | × | 36 | *Glochidion acuminatum* | × |
| 3 | *Acer palmatum* var. *pubescens* | × | 37 | *Helicia cochinchinensis* | × |
| 4 | *Adinandra formosana* var. *formosana* | × | 38 | *Helicia formosana* | × |
| 5 | *Antidesma japonicum* var. *densiflorum* | × | 39 | *Hydrangea angustipetala* | × |
| 6 | *Barthea barthei* | × | 40 | *Hydrangea aspera* | |
| 7 | *Callicarpa formosana* var. *formosana* | | 41 | *Hydrangea paniculata* | × |
| 8 | *Callicarpa randaiensis* | | 42 | *Idesia polycarpa* | |
| 9 | *Camellia brevistyla* | × | 43 | *Ilex sugerokii* var. *brevipedunculata* | × |
| 10 | *Carpinus rankanensis* | × | 44 | *Ilex ficoidea* | × |
| 11 | *Castanopsis cuspidata* var. *carlesii* | × | 45 | *Ilex formosana* | × |
| 12 | *Castanopsis uraiana* | × | 46 | *Ilex goshiensis* | × |
| 13 | *Chamaecyparis formosensis* | × | 47 | *Ilex hayatana* | × |
| 14 | *Chamaecyparis obtusa* var. *formosana* | × | 48 | *Ilex rarasanensis* | × |
| 15 | *Cinnamomum macrostemon* | × | 49 | *Ilex rotunda* | × |
| 16 | *Cinnamomum subavenium* | × | 50 | *Ilex suzukii* | |
| 17 | *Clerodendrum cyrtophyllum* | | 51 | *Ilex tugitakayamensis* | × |
| 18 | *Clerodendrum trichotomum* | | 52 | *Illicium anisatum* | × |
| 19 | *Cleyera japonica* | × | 53 | *Illicium arborescens* | × |
| 20 | *Cornus kousa* subsp. *chinensis* | × | 54 | *Itea parviflora* | × |
| 21 | *Cryptocarya chinensis* | × | 55 | *Lasianthus appressihirtus* var. *appressihirtus* | × |
| 22 | *Cunninghamia lanceolata* var. *konishii* | × | 56 | *Lasianthus appressihirtus* var. *maximus* | |
| 23 | *Daphniphyllum himalaense* subsp. *macropodum* | × | 57 | *Ligustrum liukiuense* | × |
| 24 | *Daphniphyllum glaucescens* subsp. *oldhamii* | × | 58 | *Ligustrum sinense* | |
| 25 | *Dendropanax dentiger* | × | 59 | *Lindera megaphylla* | |
| 26 | *Deutzia pulchra* | | 60 | *Lithocarpus brevicaudatus* | × |
| 27 | *Diospyros morrisiana* | × | 61 | *Lithocarpus hancei* | |
| 28 | *Elaeagnus formosana* | | 62 | *Litsea acuminata* | × |
| 29 | *Elaeocarpus japonicus* | × | 63 | *Litsea elongata var. mushaensis* | × |
| 30 | *Elaeocarpus sylvestris* var. *sylvestris* | | 64 | *Machilus japonica* var. *japonica* | × |
| 31 | *Elaeagnus thunbergii* | | 65 | *Machilus zuihoensis* var. *mushaensis* | × |
| 32 | *Eurya crenatifolia* | × | 66 | *Machilus thunbergii* | × |
| 33 | *Eurya glaberrima* | × | 67 | *Machilus zuihoensis* var. *zuihoensis* | × |
| 34 | *Eurya loquaiana* | × | 68 | *Meliosma squamulata* | × |

| Nr | Species | Traits |
|---|---|---|
| 69 | *Michelia compressa* | × |
| 70 | *Microtropis fokienensis* | |
| 71 | *Myrica rubra* | |
| 72 | *Myrsine seguinii* | × |
| 73 | *Nageia nagi* | |
| 74 | *Neolitsea aciculata* | × |
| 75 | *Neolitsea acuminatissima* | × |
| 76 | *Neolitsea konishii* | × |
| 77 | *Neolitsea sericea* var. *sericea* | × |
| 78 | *Osmanthus heterophyllus* | × |
| 79 | *Osmanthus matsumuranus* | × |
| 80 | *Phyllanthus oligospermus* | |
| 81 | *Pourthiaea beauverdiana* var. *notabilis* | × |
| 82 | *Pourthiaea villosa* var. *parvifolia* | |
| 83 | *Prunus buergeriana* | |
| 84 | *Prunus obtusata* | |
| 85 | *Prunus phaeosticta* var. *phaeosticta* | × |
| 86 | *Prunus transarisanensis* var. *takasagomontana* | × |
| 87 | *Pyrenaria shinkoensis* | × |
| 88 | *Quercus gilva* | × |
| 89 | *Quercus glauca* var. *glauca* | |
| 90 | *Quercus longinux* var. *longinux* | × |
| 91 | *Quercus morii* | × |
| 92 | *Quercus sessilifolia* | × |
| 93 | *Quercus stenophylloides* | × |
| 94 | *Rhododendron formosanum* | × |
| 95 | *Rhododendron leptosanthum* | × |
| 96 | *Rhododendron pseudochrysanthum* | × |
| 97 | *Schefflera octophylla* | × |
| 98 | *Schima superba* var. *superba* | × |
| 99 | *Skimmia reevesiana* | × |
| 100 | *Sorbus randaiensis* | × |
| 101 | *Stachyurus himalaicus* | |
| 102 | *Sycopsis sinensis* | × |
| 103 | *Symplocos arisanensis* | × |
| 104 | *Symplocos glauca* | × |
| 105 | *Symplocos konishii* | |
| 106 | *Symplocos macrostroma* | × |
| 107 | *Symplocos migoi* | × |
| 108 | *Symplocos setchuensis* | |
| 109 | *Symplocos stellaris* | × |

| Nr | Species | Traits |
|---|---|---|
| 110 | *Symplocos theophrastifolia* | × |
| 111 | *Symplocos wikstroemiifolia* | × |
| 112 | *Syzygium buxifolium* | × |
| 113 | *Ternstroemia gymnanthera* | × |
| 114 | *Tricalysia dubia* | × |
| 115 | *Trochodendron aralioides* | × |
| 116 | *Tsuga chinensis* var. *formosana* | × |
| 117 | *Viburnum erosum* | |
| 118 | *Viburnum formosanum* | |
| 119 | *Viburnum luzonicum* | |
| 120 | *Viburnum foetidum* var. *rectangulatum* | × |
| 121 | *Viburnum sympodiale* | × |
| 122 | *Viburnum urceolatum* | |

**Understory species**

| Nr | Species | Traits |
|---|---|---|
| 123 | *Acystopteris taiwaniana* | × |
| 124 | *Acystopteris tenuisecta* | |
| 125 | *Alsophila podophylla* | × |
| 126 | *Alsophila spinulosa* | |
| 127 | *Arachniodes amabilis* var. *amabilis* | × |
| 128 | *Arachniodes festina* | × |
| 129 | *Arachniodes pseudoaristata* | × |
| 130 | *Asplenium normale* var. *normale* | × |
| 131 | *Athyrium iseanum* var. *angustisectum* | |
| 132 | *Athyrium arisanense* | × |
| 133 | *Athyrium delavayi* var. *delavayi* | |
| 134 | *Athyrium nakanoi* | × |
| 135 | *Athyrium opacum* | |
| 136 | *Blechnopsis orientalis* | |
| 137 | *Cheiropleuria integrifolia* | |
| 138 | *Coniogramme intermedia* | |
| 139 | *Coryphopteris angulariloba* | |
| 140 | *Coryphopteris castanea* | × |
| 141 | *Dennstaedtia scabra* | × |
| 142 | *Deparia formosana* | × |
| 143 | *Dicranopteris tetraphylla* | |
| 144 | *Diplazium sp.* | |
| 145 | *Diplazium dilatatum* | × |
| 146 | *Diplazium doederleinii* | × |
| 147 | *Diplazium donianum* var. *donianum* | × |
| 148 | *Diplopterygium glaucum* | × |
| 149 | *Diplazium kawakamii* var. *kawakamii* | × |

| Nr | Species | Traits | Nr | Species | Traits |
|---|---|---|---|---|---|
| 150 | *Diplazium mettenianum* | × | 175 | *Microlepia hookeriana* | × |
| 151 | *Diplazium okinawaense* | | 176 | *Microlepia obtusiloba* | |
| 152 | *Diplazium petrii* | × | 177 | *Monachosorum henryi* | × |
| 153 | *Diplazium pullingeri* | × | 178 | *Nephrolepis cordifolia* | |
| 154 | *Diplazium virescens* var. *virescens* | | 179 | *Odontosoria chinensis* | |
| 155 | *Dryopteris formosana* | × | 180 | *Plagiogyria adnata* | × |
| 156 | *Dryopteris hasseltii* | × | 181 | *Plagiogyria euphlebia* | × |
| 157 | *Dryopteris hendersonii* | × | 182 | *Plagiogyria falcata* | × |
| 158 | *Dryopteris lepidopoda* | | 183 | *Plagiogyria glauca* | × |
| 159 | *Dryopteris melanocarpa* | × | 184 | *Plagiogyria stenoptera* | × |
| 160 | *Dryopteris paleolata* | × | 185 | *Polystichum hancockii* | × |
| 161 | *Dryopteris polita* | | 186 | *Polystichum integripinnum* | × |
| 162 | *Dryopteris subexaltata* | | 187 | *Polystichum parvipinnulum* | × |
| 163 | *Dryopteris subtriangularis* | × | 188 | *Pronephrium gymnopteridifrons* | × |
| 164 | *Dryopteris wuzhaohongii* | × | 189 | *Pteris bella* | × |
| 165 | *Histiopteris incisa* | × | 190 | *Pteris tokioi* | × |
| 166 | *Huperzia serrata* | × | 191 | *Pteris wallichiana* | |
| 167 | *Hymenasplenium adiantifrons* | × | 192 | *Selaginella delicatula* | |
| 168 | *Leptogramma tottoides* | | 193 | *Selaginella doederleinii* subsp. *doederleinii* | |
| 169 | *Lindsaea bonii* | × | 194 | *Selaginella labordei* | |
| 170 | *Lindsaea chienii* | × | 195 | *Selaginella remotifolia* | |
| 171 | *Lycopodiella cernua* | | 196 | *Stegnogramma griffithii* | × |
| 172 | *Metathelypteris gracilescens* | × | 197 | *Stegnogramma wilfordii* | × |
| 173 | *Metathelypteris laxa* | | 198 | *Woodwardia unigemmata* | |
| 174 | *Metathelypteris uraiensis* | × | | | |

**Appendix S4. Parameter estimates of the reduced redundancy analyses (RDA) for the species composition including all species (all) and only species for which traits were measured (subset) for overstory and understory, and for the overstory trait composition including all species (all) and excluding deciduous species (subset).** Test statistic (F) for each retained predictor and full model adjusted $R^2$ provided. For soil principal components 'soil NPK', 'soil pH' and 'soil Cu', see Table 1. $^{(*)}$0.10 ≥ p-value > 0.05; $^{*}$0.05 ≥ p-value > 0.01; $^{**}$0.01 ≥ p-value > 0.001; $^{***}$0.001 ≥ p-value. $^{sqrt}$ = square root transformation, $^{sq}$ = squared transformation. Fog = ground fog frequency.

| | climate proxies | | | soil | | | | | |
|---|---|---|---|---|---|---|---|---|---|
| | elevation | fog | heat load$^{sq}$ | soil depth | soil NPK | soil pH | soil Cu | soil rockiness$^{sqrt}$ | $R^2$ |
| **Species composition** | | | | | | | | | |
| overstory (all) | 10.4$^{***}$ | 4.7$^{***}$ | 1.6$^{(*)}$ | - | 1.7$^{*}$ | 3.2$^{***}$ | 1.6$^{(*)}$ | - | 29.2 |
| overstory (subset) | 11.4$^{***}$ | 5.1$^{***}$ | 1.6$^{(*)}$ | - | 1.8$^{*}$ | 3.2$^{***}$ | 1.7$^{(*)}$ | - | 31.1 |
| understory (all) | 10.4$^{***}$ | 7.3$^{***}$ | 1.9$^{*}$ | - | - | 3.9$^{***}$ | - | - | 29.8 |
| understory (subset) | 9.1$^{***}$ | 7.9$^{***}$ | 2.0$^{*}$ | - | - | 4.0$^{***}$ | - | - | 31.1 |
| **CM Trait composition** | | | | | | | | | |
| overstory (all) | 10.0$^{***}$ | 4.3$^{**}$ | - | - | 4.7$^{**}$ | 13.1$^{***}$ | 7.8$^{***}$ | - | 49.4 |
| overstory (subset) | 17.8$^{***}$ | 6.5$^{**}$ | - | - | 4.5$^{**}$ | 6.8$^{***}$ | 5.0$^{**}$ | - | 50.9 |

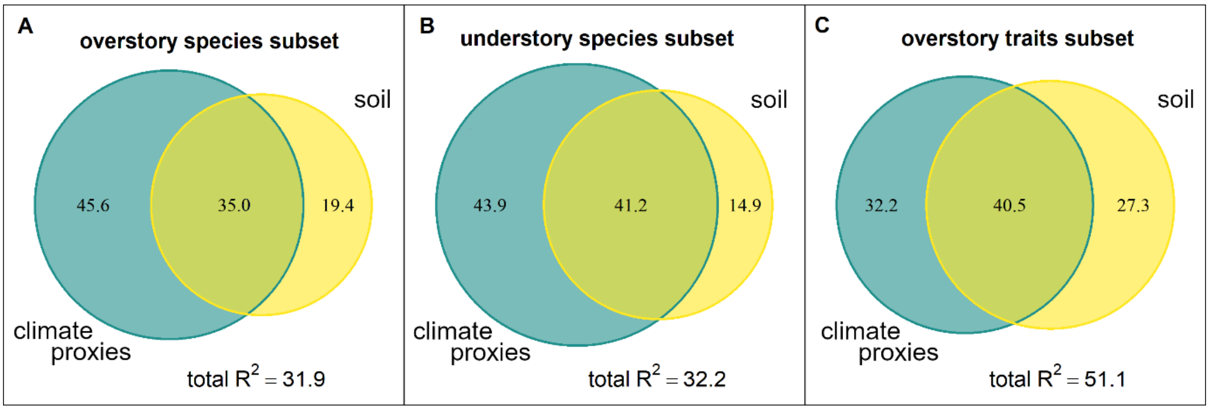


**Appendix S5. Venn diagram visualizing the variation partitioning between climate proxy and soil variable effects on A. the overstory plot × species matrix including only species for which traits were measured, B. the understory plot × species matrix including only species for which traits were measured, C. the overstory plot × community mean (CM) trait matrix excluding deciduous species, using redundancy analysis (RDA).** Numbers in the Venn diagrams correspond to the proportion of the total explained variation. The total explained variance (adjusted $R^2$) is also presented.

**Appendix S6. Separate pairwise linear regressions between understory' and overstory' community mean (CM) traits and diversity measures, for both overstory traits excluding and including deciduous species.** F statistic and p-value (after FDR correction) for each predictor, and adjusted model $r^2$ given. Note that for SLA, the model contains F and p values for the two parameters. Significant results ($p < 0.05$) in bold. $\delta^{13}C$ = the leaf $^{13}C/^{12}C$ stable isotope ratio, $\delta^{15}N$ = the leaf $^{15}N/^{14}N$ stable isotope ratio, EWT = equivalent water thickness, LDMC = leaf dry matter content, leaf N = leaf nitrogen content, SLA = specific leaf area. a = logarithmic transformation of CM trait for understory traits, b = square transformation of CM trait for overstory traits.

| | **Traits incl. deciduous sp.** | | | **Traits excl. deciduous sp.** | | |
|---|---|---|---|---|---|---|
| | **F** | **p** | **$r^2$** | **F** | **p** | **$r^2$** |
| **CM trait** | | | | | | |
| SLA | **SLA:5.3**<br>**SLA$^2$: 14.5** | **SLA: 0.016**<br>**SLA$^2$: <0.001** | **38.8** | **SLA:1.6**<br>**SLA$^2$: 6.6** | **SLA: 0.276**<br>**SLA$^2$: 0.030** | **15.0** |
| LDMC | **31.2** | **<0.001** | **34.3** | **26.1** | **<0.001** | **30.2** |
| Log leaf area | 0.7 | 0.502 | <0.1 | 0.9 | 0.908 | <0.1 |
| leaf thickness | **5.7** | **0.040** | **7.4** | **17.6** | **0.001** | **22.3** |
| leaf chlorophyll content | 0.6 | 0.849 | <0.1 | 0.1 | 0.849 | <0.1 |
| EWT | 2.5 | 0.181 | 2.6 | 0.3 | 0.682 | <0.1 |
| $\delta^{13}C$ | 4.9 | 0.055 | 6.3 | 2.1 | 0.223 | 1.8 |
| $\delta^{15}N^{ab}$ | **8.4** | **0.016** | **11.3** | **5.9** | **0.039** | **7.7** |
| leaf N | **15.4** | **0.001** | **19.9** | **7.6** | **0.020** | **10.2** |
| **Diversity** | | | | | | |
| species richness | 1.6 | 0.276 | <0.1 | - | - | - |
| functional dispersion | **16.3** | **0.001** | **21.7** | - | - | - |
| functional equability | 4.2 | 0.075 | 5.7 | - | - | - |

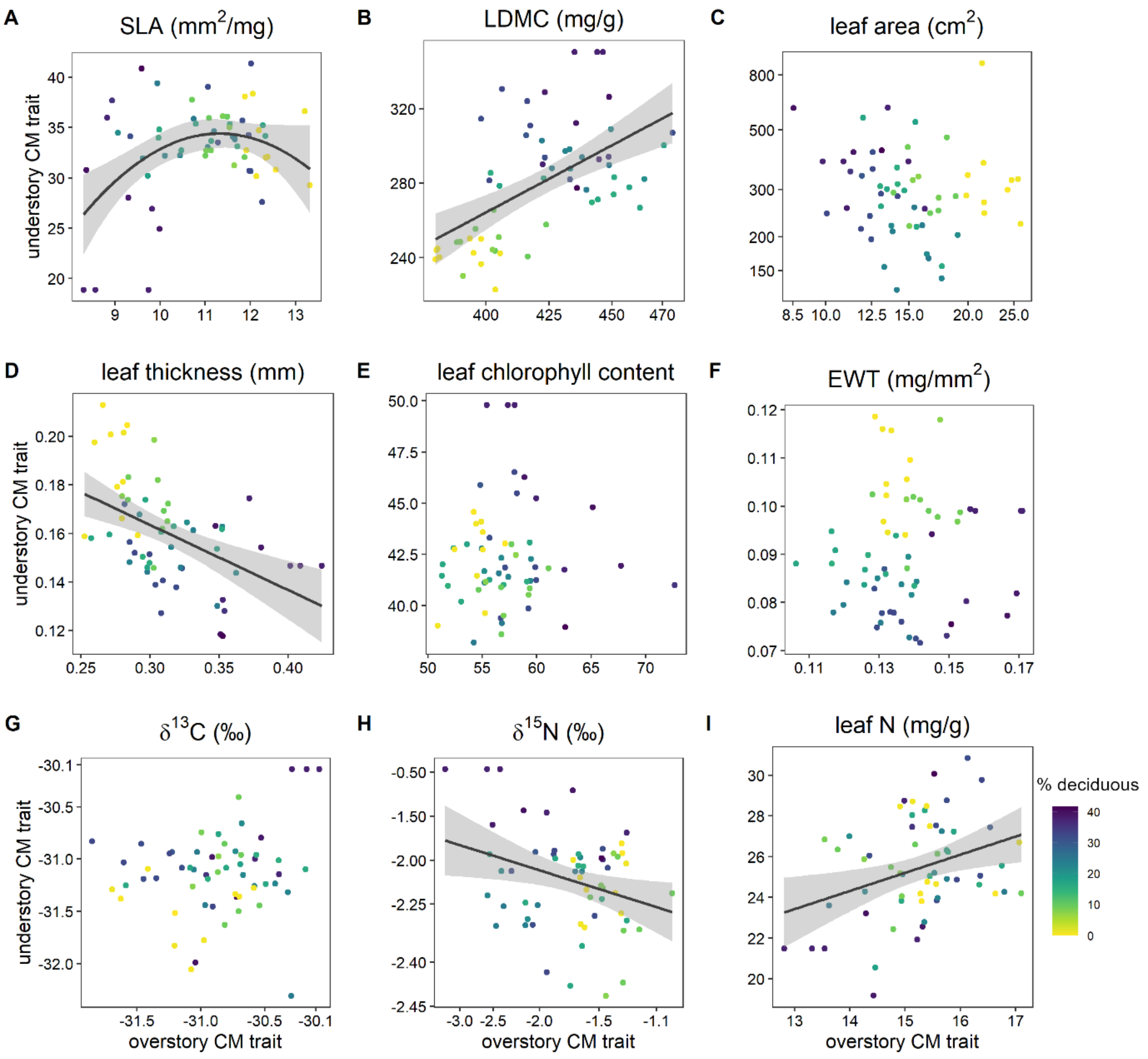


**Appendix S7. Scatterplots for pairwise regressions between plot-level overstory and understory community mean (CM) trait values; for overstory excluding deciduous species.** Regression line + SE presented for significant regressions (see Appendix S6). Each datapoint corresponds to one vegetation plot, with colors indicating plot elevation. $\delta^{13}C$ = the leaf $^{13}C/^{12}C$ stable isotope ratio, $\delta^{15}N$ = the leaf $^{15}N/^{14}N$ stable isotope ratio, EWT = equivalent water thickness, Lth = leaf thickness.

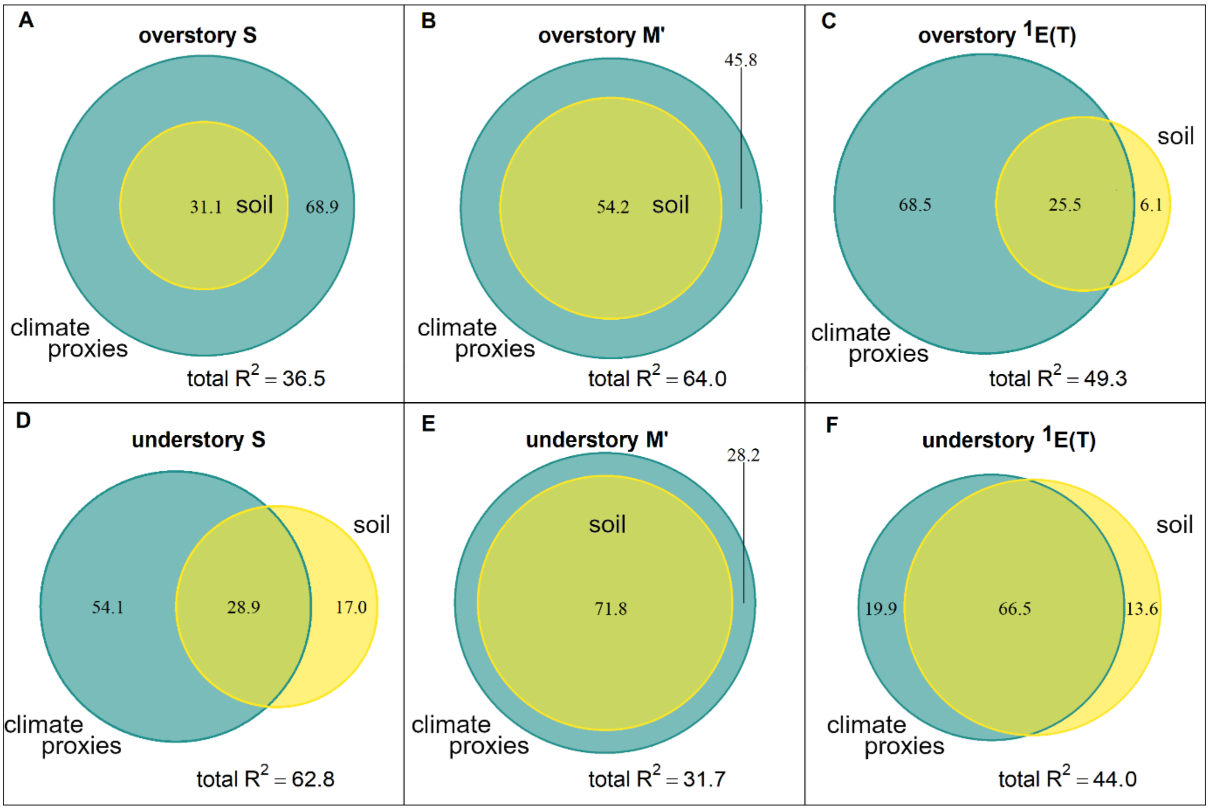


**Appendix S8. Venn diagrams visualizing the variation partitioning between climate proxy and soil variable effects on over- and understory species richness (S), functional divergence (M') and functional equability ($^{1}E(T)$), using generalized linear models (GLM).** Numbers in the Venn diagrams correspond to the proportion of the total explained variation. The total explained variance (adjusted $R^2$) is also presented.

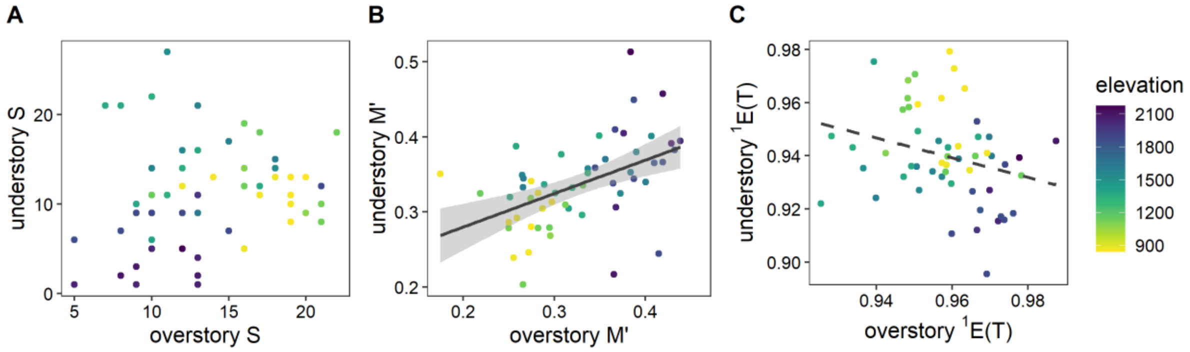


**Appendix S9. Scatterplots for pairwise regressions between plot-level overstory and understory A. species richness (S), B. functional dispersion (M') and C. functional equability ($^1$E(T)).** Regression line + SE presented for significant regressions, dashed line for marginally significant regression (see Appendix S6). Each datapoint corresponds to one vegetation plot, with colors indicating plot elevation.
